# Matrix confinement modulates 3D spheroid sorting and burst-like collective migration

**DOI:** 10.1101/2023.07.23.549940

**Authors:** Grace Cai, Xinzhi Li, Shan-Shan Lin, Samuel J. Chen, Nicole C. Rodgers, Katherine M. Koning, Dapeng Bi, Allen P. Liu

## Abstract

While it is known that cells with differential adhesion tend to segregate and preferentially sort, the physical forces governing sorting and invasion in heterogeneous tumors remain poorly understood. To investigate this, we tune matrix confinement, mimicking changes in the stiffness and confinement of the tumor microenvironment, to explore how physical confinement influences individual and collective cell migration in 3D spheroids. High levels of confinement lead to cell sorting while reducing matrix confinement triggers the collective fluidization of cell motion. Cell sorting, which depends on cell-cell adhesion, is crucial to this phenomenon. Burst-like migration does not occur for spheroids that have not undergone sorting, regardless of the degree of matrix confinement. Using computational Self-Propelled Voronoi modeling, we show that spheroid sorting and invasion into the matrix depend on the balance between cell-generated forces and matrix resistance. The findings support a model where matrix confinement modulates 3D spheroid sorting and unjamming in an adhesion-dependent manner, providing insights into the mechanisms of cell sorting and migration in the primary tumor and toward distant metastatic sites.

## Introduction

The primary tumor microenvironment plays a crucial role in facilitating the movement of cancer cells beyond the primary tumor. These cells navigate through the stroma, eventually infiltrating nearby microvessels through the process of intravasation [1–3]. In confined tumors that exhibit a high cellular density, cancer cells must proliferate and migrate in order to promote tumor growth and metastasis. However, unchecked proliferation within these restricted spaces can lead to confinement stress [4–6]. This stress in turn induces changes in cell-cell interactions and influences tumor development and compartmentalization [7,8]. Many cell-cell interactions depend on E-cadherin-based cell-cell junctions to preserve tissue cohesion and maintain active force transmission [9–11]. During cancer metastasis, E-cadherin is downregulated in epithelial-to-mesenchymal transition (EMT), which destabilizes cell-cell junctions and promotes invasiveness [12,13]. Cancer invasion and metastasis have been shown to occur in both E-cadherin-expressing and -deficient tumors [14]. However, the impact of the mechanical properties of the tumor microenvironment on intercellular interactions, especially in heterogeneous tumors, remains unclear.

A heterogeneous cell composition effectively replicates the physiological tumor microenvironment and introduces heterotypic cell-cell interactions [15,16]. Mixtures of two cell types with differences in cell-cell adhesion have been observed to undergo sorting [17,18], however the cell-cell and cell-substrate interactions driving this phenomenon in tumors are not well understood. Recent work suggests that changes to cell-cell adhesion associated with EMT are related to cellular jamming and unjamming [19,20]. When cells are jammed, they behave as a solid-like state in which individual cellular motion becomes arrested, and unjamming represents a transition to a fluid-like state in which cells exchange neighbors and the tissue flows in response to fluctuations [21,22]. In essence, the physical mechanisms that orchestrate jamming— unjamming behavior are driven by cellular properties, such as cell-cell and cell-substrate interactions [23].

Migrating cells within the tumor microenvironment encounter various physical cues that alter their behavior and migration patterns, including ECM stiffness, fluid shear stress, interstitial fluid pressure, and fluid viscosity [24–27]. In particular, microenvironmental conditions involving cell-substrate interactions play a pivotal role in regulating 3D collective migration [24,28]. Many cell types rely on integrin-based traction forces to facilitate migration, and enhanced cell-substrate adhesion is associated with increased cell spreading and greater motility [23,29,30]. Notably, the mechanical properties of the substrate itself are also a significant factor in influencing cell behavior, and cells tend to exhibit more pronounced spreading and faster migration on stiffer substrates [31–33].

Collagen gels provide a versatile method for adjusting substrate stiffness, achieved through measures such as increasing collagen concentration or employing more potent crosslinking agents [34–36]. Hybrid hydrogels, such as GelMA-collagen gels, offer a precise mechanism for finely tuning both mechanical properties and ligand density [37]. Similarly, collagen-alginate gels allow for the independent modulation of the stiffness of the alginate component, with collagen introduction regulating ligand density for cell adhesion and interactions. Collagen-alginate hydrogels display viscoelastic properties during gelation; post-curing, they behave more like solids than viscous fluids [38]. Changing the calcium concentration does not significantly alter the distribution of their pore sizes [38–40]. Moreover, varying the degree of calcium crosslinking does not impact the availability of cell adhesion motifs on the collagen network [38]. By leveraging calcium and calcium chelators such as ethylene glycol tetraacetic acid (EGTA), we can manipulate the hydrogel’s stiffness in a time-dependent manner, mirroring the physical properties observed in the tumor microenvironment. Invasive cancer cells possess the capability to degrade collagen through the secretion of matrix metalloproteinases (MMPs) [41]. Thus, alginate finds greater relevance in contexts where degradation is limited or not feasible. In our study, we aim to investigate the specific influence of matrix stiffness on cell sorting and invasion within 3D spheroids.

In our investigation into the impact of matrix confinement on cancer migration, we propose that tumor cell migration occurs in two distinct stages, in which cell sorting leads to pressure-driven cellular invasion. In this study, we tune matrix confinement to illustrate the critical role that ECM stiffness plays in tumor development and distant metastasis. Increased matrix confinement triggers cell sorting within a spheroid, causing cells sorted to the spheroid core to become jammed. A reduction in matrix confinement causes the collective fluidization of cell motion, propelling normal epithelial cells and cancer cells into the matrix with high velocity. Using a computational Self-Propelled Voronoi (SPV) model to simulate a heterogeneous tissue, we confirm the experimental finding and further show that decreasing the cell-medium contact tension downregulates confinement stress and leads to cell invasion. The results yield insights into the interplay between confinement stress, cell-cell adhesion, and 3D jamming–unjamming transitions in breast cancer metastasis.

## Materials and Methods

### Cell culture

The cell lines MCF10A, MCF7 and MDA-MB-231 were gifts from Sofia Merajver (University of Michigan). MCF10A cells were obtained from Gloria Heppner at the Michigan Cancer Foundation where the cell line was originally developed, MCF7 cells were originally obtained from ATCC, and MDA-MB-231 cells were obtained from Janet Price (MD Anderson Cancer Center) where the cell line was originally developed. MCF10A cells were cultured in DMEM/F12 medium (Gibco) supplemented with 5% horse serum, 20 ng/ml epidermal growth factor (EGF), 0.5 μg/ml hydrocortisone, 100 ng/ml cholera toxin and 10 μg/ml insulin. MCF7 and MDA-MB-231 cells were cultured in DMEM medium (Gibco) supplemented with 10% FBS. Cells were cultured in a humidified atmosphere containing 5% CO_2_ at 37°C.

### Stable cell lines

Stable cell lines expressing EGFP or Lifeact-RFP were generated via lentiviral transduction. EGFP lentivirus was produced from pSMPVW-EGFP obtained from Andrew Tai (University of Michigan). The lentiviral transfer plasmid pLVX-puro-RFP-Lifeact was cloned from RFP-Lifeact obtained from Gaudenz Danuser (UT-Southwestern). Lentiviruses were generated by transfecting HEK 293T cells with the transfer vector, psPAX2 packaging vector, and pMD2.G envelope vector. Viral supernatant was collected 48 h after transfection and used to infect the target cell lines. After 24 h, transduced cells were selected with 2 μg/mL puromycin for 5 days.

### E-cadherin knockdown

MCF10A cells expressing inducible shRNA knockdown of E-cadherin was generated using a transfer plasmid provided by Valerie Weaver at UCSF [42]. The transfer vector consisted of a modified pLKO.1 neo plasmid (Addgene) with expression of the shRNA sequences under control of 3× copies of the lac operator. The E-cadherin shRNA had the following sequence: 5’ - GAACGAGGCTAACGTCGTAAT - 3’; scramble shRNA (Sigma #SHC002) had the following sequence: CCGGCAACAAGATGAAGAGCACCAACTCGAGTTGGTGCTCTTCATCTTGTTGTTTTT. MCF10A cells were transduced with E-cadherin shRNA or scramble non-targeting control shRNA for 48 h. shCDH1 and scramble cells were selected with 200 μg/ml G-418 (Sigma) and 2 μg/mL puromycin, respectively. E-cadherin knockdown was induced in shCDH1 cells by adding 200 μM isopropyl-β-D-thiogalactoside (IPTG; Sigma) 72 h prior to experiments.

### Encapsulation of tumor spheroids in a collagen-alginate hydrogel

Inverse pyramidal PDMS microwells (AggreWell™, Stem Cell Technologies) were treated with anti-adherence rinsing solution (Stem Cell Technologies) to prevent cell adhesion. Cells were detached with 0.25% trypsin-EDTA (Life Technologies) and added to the microwells at a concentration of ∼1,000 cells per microwell. For co-culture spheroids, MCF10A cells were mixed with MDA-MB-231, MCF7 or MCF10A shCDH1 KD cells at a 2:1 ratio. The cells were centrifuged at 300g for 5 min to aggregate the cells at the bottom of the microwells, and spheroids formed overnight. The following day, the spheroids were harvested from the microwells and encapsulated in collagen-alginate hydrogels consisting of 3 mg/ml type I rat tail collagen (RatCol, Advanced BioMatrix) and 0.25% alginate (Nalgin HG, Tilley Chemical). Subsequently, the hydrogels were incubated at 37°C for 1 h for collagen polymerization, and then the encapsulated spheroids were imaged (day 0). Alginate crosslinking density and hydrogel stiffness were independently modulated through the concentration of CaCl_2_ that was added to the cell culture medium after imaging on day 0. On day 4, the spheroids were imaged and then washed 1× with PBS. Afterward, the spheroids were either fixed for immunofluorescence staining or incubated with 0, 5 or 10 mM EGTA in PBS for 1 h to reverse the alginate crosslinking process. The solution was then replaced with fresh medium, and on day 6, the spheroids were fixed for immunofluorescence staining.

FITC-labeled collagen-alginate hydrogels were prepared by incubating 2.5% FITC-labeled rat collagen on ice overnight. The following day, MCF10A and MCF7 Lifeact-RFP co-culture spheroids were harvested from the microwells and encapsulated in collagen-alginate hydrogels consisting of 2.5% FITC-labeled 2.5 mg/ml type I rat tail collagen and 0.25% alginate treated with different amounts of CaCl_2_.

The Young’s moduli of collagen-alginate hydrogels were measured after 4 days of incubation with 0, 5 or 10 mM CaCl_2_ or after treatment with 0, 5, or 10 mM EGTA in PBS for 1 h following 4 days of incubation in 10 mM CaCl_2._ Measurements were made using a MicroSquisher (CellScale). A microbeam with a diameter of 203.2 μm, modulus of 411,000 MPa, and length of 59.5 mm was fixed to a 0.75 mm diameter glass bead and mounted to the vertical actuator. The samples were submerged in a solution of 1% BSA in PBS to reduce adhesion and compressed 3– 4 times at different locations with a vertical displacement of 50–150 μm and at a loading rate of 1 μm/s. The Young’s modulus was calculated using the linear slope of the stress-strain curve.

Viscoelasticity characterizations were performed using a Discovery HR-2 rheometer (TA Instruments) with 20 mm top- and bottom-plate stainless steel geometries. After incubation with 0, 5, or 10 mM CaCl_2_ for 24 h, collagen-alginate hydrogels were loaded onto the center of the bottom plate, and the 20 mm flat top plate was quickly lowered to secure the gel. Prior to loading, sandpaper was glued onto the top and bottom plate to prevent hydrogel slipping. A time sweep was conducted at 1 rad/s and 1% strain for 1,000 seconds, and measurements were taken every 10 seconds. The loss tangent for each measurement was calculated by the TRIOS software (TA Instruments), and the averages of each run were recorded.

### Confocal microscopy

Images were taken using an oil immersion UplanFL N 10 x/1.30 NA (Olympus) objective on an inverted microscope (Olympus IX-81) equipped with an iXON3 EMCCD camera (Andor Technology), National Instrument DAQ-MX controlled laser (Solamere Technology), and a Yokogawa CSU-X1 spinning disk confocal unit. Z-stack images of spheroids expressing EGFP or Lifeact-RFP and fluorescently labeled for DAPI were taken at excitation wavelengths of 488, 561 and 405 nm, respectively. Z-stack images of spheroids fluorescently labeled for E-cadherin, vimentin, or F-actin (by 670-phalloidin) were taken at an excitation wavelength of 640 nm.

### Immunofluorescence staining and image analysis

Spheroids embedded in hydrogels were washed with PBS and fixed with 4% paraformaldehyde for 1 h, washed with PBS, and permeabilized with 0.1% Triton X-100 in PBS for 4 h. Then, cells were washed with PBS and blocked with 3% BSA in PBS overnight at 4°C. The following morning, cells were incubated with a mouse anti-E-cadherin antibody at 1:500 (610181, BD Biosciences) or a rabbit anti-vimentin antibody at 1:400 (D21H3, Cell Signaling) in 3% BSA overnight at 4°C. Next, cells were washed 3× with PBS for 30 minutes per wash and incubated with DAPI and a secondary antibody in 3% BSA overnight at 4°C. Hydrogels were washed 3× with PBS as described above and imaged by spinning disk confocal microscopy.

To quantify cell sorting and the E-cadherin signal, the median slice from each z-stack was analyzed. Using the Fit Ellipse and Centroid options in Fiji, the coordinates of the center of the spheroid and the semi-major axis of the spheroid were extracted. Using the Multi-point tool, the coordinates of individual cells were recorded. As a measure of cell sorting, the distance index (DI) was calculated by dividing the cell’s distance from the spheroid center by the spheroid’s semi-major axis [18]. The E-cadherin signal at cell–cell contacts was quantified by measuring fluorescence intensity at the cell membrane and was calculated per cell by dividing the integrated density by the number of cells (indicated by DAPI) for each image.

ImageJ was used to generate maximum projection images of FITC-labeled collagen gel and fluorescence intensity line scans. Line scans (3-pixel width) of about 350 μm were drawn across the center of the spheroid. Fluorescence intensity was measured along the line and background subtraction was performed by subtracting the minimum intensity along the scan from all measurements. “Fire” LUT was used for image color representation.

### Time-lapse imaging and analysis

For time-lapse imaging, images were acquired on an Olympus IX-81 inverted microscope, as previously described, or a Nikon-A1 laser scanning confocal microscope, equipped with an environmental chamber. Cell motility was recorded at 1-hour intervals over 18 or 24 h, with a 4 μm z-step. Individual cell trajectories were obtained using TrackMate in Fiji, where the LoG detector was used for spot detection with median filtering and subpixel localization, and the linear motion LAP tracker was used to link spots. After exporting the tracks as XML files, cell motility rates were calculated for each spheroid [43], and MSDs were analyzed using a MATLAB per-value class for MSD analysis of particle trajectories [44] and plotted in log-log scale, where the slope gives the diffusion coefficient α.

To calculate distance index from time-lapse sequences, z-stacks captured at 0, 8, 16 and 24 h were analyzed. Using the 3D projections of z-stacks taken at 0 h, the initial time point, the centroid location and semi-major axis of each spheroid were extracted in MATLAB via the “regionprops” function, which approximates spheroids as ellipses. Next, individual cell positions were acquired in TrackMate using the median slice of each z-stack. After exporting the cell coordinates as XML files, the distance index for each cell at all four time points was calculated using custom MATLAB code.

### Western blotting

Samples were run on SDS–PAGE 4–20% Bio-Rad gels (15 well/15 μl). SDS-PAGE gels were run at a constant 120 V for 90 min. Proteins were transferred to a nitrocellulose membrane using the iBlot transfer system and the membrane was blocked in 5% milk in TBS-T for 1 h at room temperature. Blots were incubated with a primary rabbit GAPDH antibody at 1:1000 (D16H11, Cell Signaling), a mouse anti-E-cadherin antibody at 1:2000 (610181, BD Biosciences), and a primary rabbit anti-vimentin antibody at 1:1000 (D21H3, Cell Signaling) in 5% BSA in TBS-T overnight at 4 °C. GAPDH was used as a loading control for quantifying relative gene expression. Blots were washed 3× with TBS-T, incubated with secondary antibodies for 1 h at room temperature, and then washed again with TBS-T as described above. Western blots were imaged using an Azure Biosystems Sapphire System.

Western blot results were acquired for unencapsulated spheroids. In the case of spheroids encapsulated in crosslinked collagen-alginate hydrogels, the hydrogels did not break down in lysis buffer. This lack of breakdown caused the spheroids to become trapped within hydrogel fragments, leading to a very low protein yield.

### Statistical analysis

Statistical analysis was carried out in Origin and performed with one-way ANOVA followed by Tukey post-hoc multiple comparisons test. Results were collected from three independent experiments and data from individual cells or spheroids were plotted as mean ± S.E. or shown as boxplots, depending on the experiment. Statistical significance was denoted by asterisks in the figure panels, with * = *p* < 0.05, ** = *p* < 0.01, *** = *p* < 0.001.

### Self-Propelled Voronoi (SPV) model of a heterogeneous tissue

To investigate how cells behave invasively under different confinement levels, we use the recently developed SPV model [45,46]. In the SPV model, the basic degrees of freedom are the set of 2D cell centers {*r*_*i*_} and cell shapes are given by the resulting Voronoi tessellation. The complex biomechanics that govern intracellular and intercellular interactions can be coarse-grained [47–53] and expressed in terms of a mechanical energy functional for individual cell shapes.

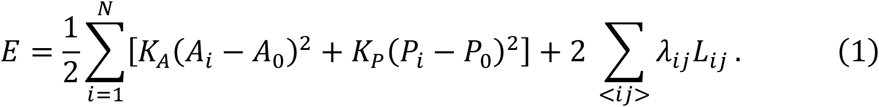

The SPV energy functional is quadratic in both cell areas *A*_*i*_ with modulus *K*_*A*_ and cell perimeters (*P*_*i*_) *P*_*i*_ with modulus *K*_*P*_. The parameters *A*_0_ and *P*_0_ set the preferred values for area and perimeter, respectively. To simulate a heterogeneous tissue [54], we have a linear tension term in the energy function Eq. (1). *λ*_*ij*_ is the junctional tension shared by cells *i* and *j* with contact edge length *L*_*ij*_. In a heterogeneous tissue with two types of cells, *λ*_*ij*_ is determined by the cell type of *i* and *j*. For example, in a tissue with two cell types A and B, we can define tensions as τ_*AA*_, τ_*BB*_ and τ_*AB*_ for A-A, B-B and A-B cell contacts.

The deformation of the actin-myosin cortex concentrated near the cell membrane is mainly responsible for changes to cell perimeters. After expanding Eq. (1), the term 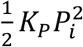 corresponds to the elastic energy associated with deforming the cortex. The linear term in cell perimeter, −*K*_*P*_*P*_0_*P*_*i*_, and *λ*_*ij*_*L*_*ij*_ represent the effective line tension in the cortex and gives rise to a ‘preferred perimeter’ *P*_0_. The value of *P*_0_ can be decreased by upregulating the contractile tension in the cortex [49,50,53] and it can be increased by upregulating cell-cell adhesion. *A*_0_ is set to be equal to the average area per cell and 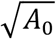 is used as the unit of length. After non-dimensionalizing Eq. (1) by 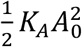 as the unit energy scale, we choose *K*_*P*_/(*K*_*A*_*A*_0_) = 1 such that the perimeter and area terms contribute equally to the cell shapes. The choice of *K*_*P*_ does not affect the results presented. The preferred cell perimeter is rescaled 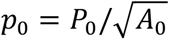 and varied between 3.7 (corresponding to the perimeter of a regular hexagon with unit area) and 4.6 (corresponding to the perimeter of an equilateral triangle with unit area) [53]. The ground states of Eq. 1 are amorphous tilings where the cells have approximately equal area, but varying perimeters as dictated by the preferred cell perimeter *p*_0_. It has been shown that at a critical value of 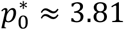, the tissue collectively undergoes a solid-fluid transition [53]. When 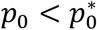, the cells must overcome finite energy barriers to rearrange and the tissue behaves as a solid, while above 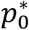, the tissue behaves as a fluid with a vanishing shear modulus as well as vanishing energy barriers for rearrangements [53].

The effective mechanical interaction force experienced by cell *i* is defined as ***F***_*i*_ = −**∇**_*i*_*E*. In addition to ***F***_*i*_, cells can also move due to self-propelled motility. Just as in SPP models, we assign a polarity vector 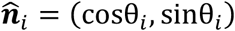 to each cell; along 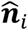 the cell exerts a self-propulsion force with constant magnitude *v*_0_/*μ*, where *μ* is the mobility (the inverse of a frictional drag). Together these forces control the over-damped equation of motion of the cell center ***r***_*i*_

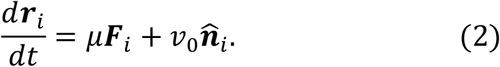

The polarity is a minimal representation of the front/rear characterization of a motile cell [55]. While the precise mechanism for polarization in cell motility is an area of intense study, here we model its dynamics as a unit vector that undergoes random rotational diffusion,

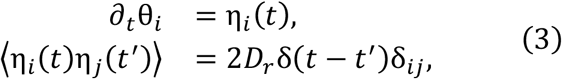

where θ_*i*_ is the polarity angle that defines 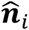, and η_*i*_(*t*) is a white noise process with zero mean and variance 2*D*_*r*_. The value of angular noise *D*_*r*_ determines the memory of stochastic noise in the system, giving rise to a persistence time scale 1/*D*_*r*_ for the polarization vector 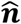.

The mechanical state of cell *i* is characterized by a local stress tensor 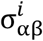 given by [56,57]

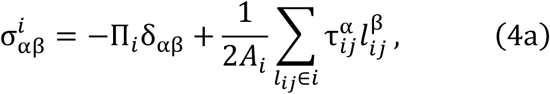

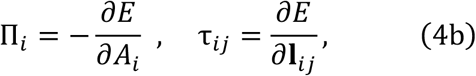

where 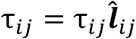 is the edge tension shared by cell *i* and *j* with 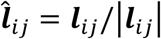, and Π_*i*_ is the hydrostatic cellular pressure. Here *i, j*, … are cell labels, and α, β denote Cartesian components. Both the tension along the edge and the intracellular pressure force perpendicularly to an edge contribute to the mechanical force balance at every vertex for a solid tissue [58]. In our simulations, the instantaneous tensions and pressures can be calculated based on Eq. (4b),

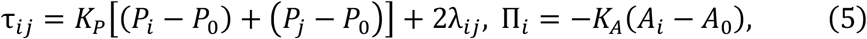

where *P*_*i*_ is the perimeter and *A*_*i*_ is the area of cell *i* respectively. The tissue stress is related to the cellular stress as

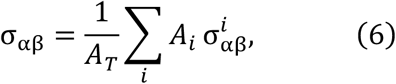

where *A*_*T*_ is the area of the tissue. The tissue compressive stress is the trace of the stress tensor *σ*_*n*_ = (*σ*_*xx*_ + *σ*_*yy*_) /2.

We simulate tissues containing two cell types (A and B) under periodic boundary condition. Each cell type has *N*_*A*_ = *N*_*B*_ = 72 cells. To initialize the simulation, a set of random cell centers are generated. Each cell is also randomly assigned a cell index and cell type label, A or B. *A*_0_ = 1, *P*_0_ = 4.2 and motility *v*_0_ = 0.3 are set to be constant values for all cells throughout the simulations. The cell shapes are obtained via Voronoi tessellations based on cell centers at each time step of the simulation. Therefore, the initial configuration is an amorphous tissue with cells A and B randomly mixed.

Junctional tensions are defined based on the types of neighboring cells, τ_*AA*_, τ_*BB*_ and τ_*AB*_ for A-A, B-B, and A-B cell contacts. In accordance with prior studies of cell sorting and the differential adhesion hypothesis [59,60], we set 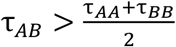 to obtain an engulfed.

Subsequently, we used the engulfed configuration as the starting state and manipulate the contact tensions between cells and medium τ_*AM*_ and τ_*BM*_, to investigate the effects of confinement on cell invasion. We solved eq. (2) using Euler’s method by running 2 *x* 10^5^ steps with time interval Δ*t* = 0.01. To quantify the cell invasion, we calculated the sorting index *I*_*A*_ = 1 − *N*_*A cluster*_/*N*_*A*_ for A cells and *I*_*B*_ = 1 *− N*_*B cluster*_/*N*_*B*_ for B cells under changing confinements. Here, *N*_*A cluster*_means the total number of isolated clusters of A cells. When the state is well sorted, there is only one A or B cell cluster, and the sorting index is *I* ≈ 1. At low confinement, cells are invading into the medium to form a mixed configuration, therefore the sorting index is *I* ≈ 0.

## Results

### 3D matrix confinement inhibits single-cell cancer migration

Spheroids are extensively utilized as a 3D *in vitro* multicellular model, mirroring the structure and function of tissues [61,62]. To form co-culture spheroids, we selected MCF10A, a non-tumorigenic breast epithelial cell line, and MDA-MB-231, a triple negative, highly invasive cancer cell line [63]. MCF10A cells have stable adherens junctions with high E-cadherin expression, whereas MDA-MB-231 cells lack E-cadherin [64]. We generated stable cell lines expressing Lifeact or a soluble fluorescence protein as markers to denote different cell types and mixed normal epithelial cells and cancer cells to replicate intratumor heterogeneity. Subsequently, we encapsulated mixed spheroids in collagen-alginate gels and manipulated the matrix stiffness by varying the concentration of CaCl_2_ to crosslink alginate (**Fig. 1A**), yielding Young’s moduli ranging from ∼0.7 kPa to ∼4.3 kPa (**Fig. 1B**), resulting in matrix confinement. This falls within the ECM stiffness range of 2-20 kPa found in breast tumors [65,66]. The crosslinking, and thereby hydrogel stiffness, was reversible by adding EGTA, a calcium chelator. By maintaining the concentration of collagen, the number of cell adhesion sites, which depend on the density of collagen fibers, remains constant.

**Figure 1.**
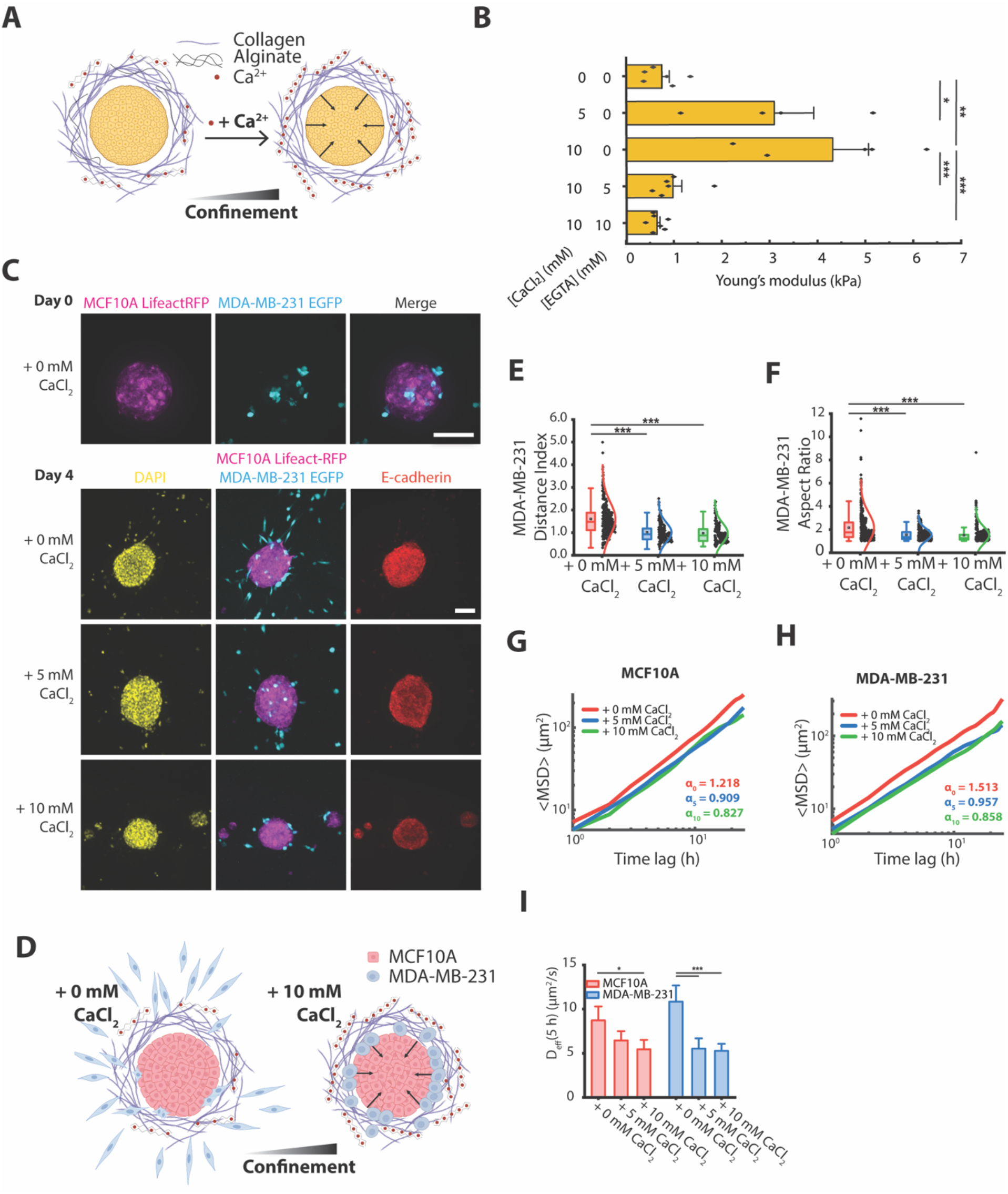
3D matrix confinement inhibits the migration of cancer cells in co-culture spheroids. **(A)** Schematic depicting a spheroid encapsulated in a collagen-alginate hydrogel. Adding Ca^2+^ to crosslink sodium alginate into calcium alginate increases matrix stiffness, leading to high matrix confinement. **(B)** Young’s moduli of collagen-alginate hydrogels following 4 days of incubation with 0, 5 or 10 mM CaCl_2_ or after a 1 h-treatment with 0, 5, or 10 mM EGTA following 4 days of incubation in 10 mM CaCl_2_. Error bars denote S.E. (**C**) Representative fluorescence images of MCF10A and MDA-MB-231 co-culture spheroids encapsulated in collagen-alginate hydrogels, imaged on days 0 and 4, and immunostained for E-cadherin and DAPI. On day 0, the spheroids were encapsulated and 0, 5 or 10 mM CaCl_2_ was added. Fluorescence images show maximum projection of the z-slices. **(D)** Schematic depicting a MCF10A (red) and MDA-MB-231 (blue) co-culture spheroid embedded in a collagen-alginate hydrogel. Addition of Ca^2+^ increases confinement stress. **(E)** Boxplot of distance index for MDA-MB-231 cells in co-culture spheroids (with MCF10A cells) after 4 days of culture. Only the median slice of the z-stack was utilized to calculate the distance index. **(F)** Boxplot of cell aspect ratio for MDA-MB-231 cells on day 4. **(G & H)** Mean square displacements (MSDs) of **(G)** MCF10A EGFP cells or **(H)** MDA-MB-231 EGFP cells in co-culture spheroids (with MCF10A Lifeact-RFP cells) plotted over an 18-h period. Each line represents the mean MSD for *n =* 12 spheroids. The plots are shown in log-log scale, and the power law exponent α is shown for each condition. **(I)** Effective diffusion coefficient (*D*_*eff*_) is shown for MCF10A and MDA-MB-231 cells for *n =* 12 spheroids per condition. Z-stacks were used to calculate MSD and *D*_*eff*_. Scale bars are 90 μm. * = *p* < 0.05, ** = *p* < 0.01, *** = *p* < 0.001.

Over 4 days, MDA-MB-231 cells sorted to the spheroid boundary and migrated as single cells into the surrounding matrix, depending on the hydrogel stiffness (**Fig. 1, C and D**). Under low confinement, single cancer cells escaped the tumor spheroid and invaded into the matrix after 4 days of culture, whereas higher confinement inhibited cancer cell migration (**Fig. 1C**). Breast carcinoma cells, which are known to be mechanically soft compared to normal cells [64], possessed low F-actin (**Fig. S1**), suggesting higher deformability than the MCF10A cells confined to the spheroid. As a measure of cell sorting and migration, a distance index was calculated for each cell using the relative distance of the cell to the spheroid center [18]. In hydrogels with low matrix stiffness, cancer cells exhibited a higher distance index (**Fig. 1E**) and were more elongated (**Fig. 1F**). Although MDA-MB-231 cells localized to the spheroid boundary after 4 days of culture regardless of matrix confinement, high confinement restricted cancer cells to the periphery of the spheroid and prevented invasion into the matrix. Furthermore, these cancer cells displayed a round morphology (**Fig. 1C**). Cancer cell invasion was not impeded in mixed spheroids cultured in collagen gels with 10 mM CaCl_2_ (**Fig. S2**), showing that the behavior observed was not caused by the presence of calcium.

To study the kinematics of 3D cancer cell migration in co-culture spheroids, we performed experiments with varying degrees of alginate crosslinking and computed the mean square displacement (MSD) in log-log scale of cells in different conditions of matrix confinement, with the average slopes (α values) representing super-diffusive (α > 1), diffusive (α = 1), and sub-diffusive (α < 1) cell motility [18,44]. Under low confinement, MDA-MB-231 cells were more diffusive than MCF10A cells (i.e., α of ∼1.51 vs. ∼1.21), whereas diffusivity was similar for both cell types under high confinement (i.e., α of ∼0.86 vs. ∼0.83, **Fig. 1, G and H**). Normal breast epithelial cells and cancer cells both demonstrated lower diffusivity when cultured in crosslinked hydrogels (**Fig. 1I**). In gels with low matrix confinement, cancer cells were more elongated, which possibly correlated with super-diffusive behavior (**Fig. 1H**), linking cell shape to diffusive motion [67,68].

### High ECM confinement drives spheroid sorting

To investigate how the metastatic potential of the cancer cell line impacts migration behavior under confinement, we generated MCF10A and MCF7 co-culture spheroids. MCF7 is a poorly metastatic breast epithelial cell line that does not express membrane type 1-matrix metalloproteinase 1 (MT1-MMP)/MMP14 [14]. As a result, MCF7 cells are unable to degrade or remodel ECM and are non-invasive. When subjected to matrix confinement, these mixed spheroids undergo different degrees of sorting, showing minimal sorting in 0 mM CaCl_2_, intermediate sorting in 5 mM CaCl_2_, and complete sorting in 10 mM CaCl_2_ (**Fig. 2, A and B**). After 4 days of culture, mixed spheroids cultured in low-stiffness gels were significantly larger than those in crosslinked gels (**Fig. S3A**), despite containing a similar number of cells. This indicates higher cell density in conditions of increased confinement (**Fig. S3B**). To quantify the spatial organization of MCF10A and MCF7 cells within spheroids as matrix confinement increased, we measured the distance index for the subpopulations, which yielded notable cell sorting under high confinement (**Fig. 2C**). Cell sorting was evident, as the subpopulations were distinctly separated into the core and edge regions of the spheroids **(Fig. 2C and Movie S1)**. This observation is correlated with differences in E-cadherin expression, as measured by fluorescence intensity normalized to the number of each cell type, where MCF10A cells at the core expressed higher E-cadherin levels than MCF7 cells localized to the periphery (**Fig. 2D**). As a measure of the differential E-cadherin expression in sorted spheroids, the relative adhesion ratio was calculated for individual spheroids as the ratio of E-cadherin expression of the adhesive cell type to the less adhesive cell type, based on their fluorescence intensity (**Fig. 2E**). The adhesion ratio increased with spheroid sorting under high matrix confinement, indicating that sorting is indeed linked to intercellular adhesion strength, consistent with the differential adhesion hypothesis (DAH) [59].

**Figure 2.**
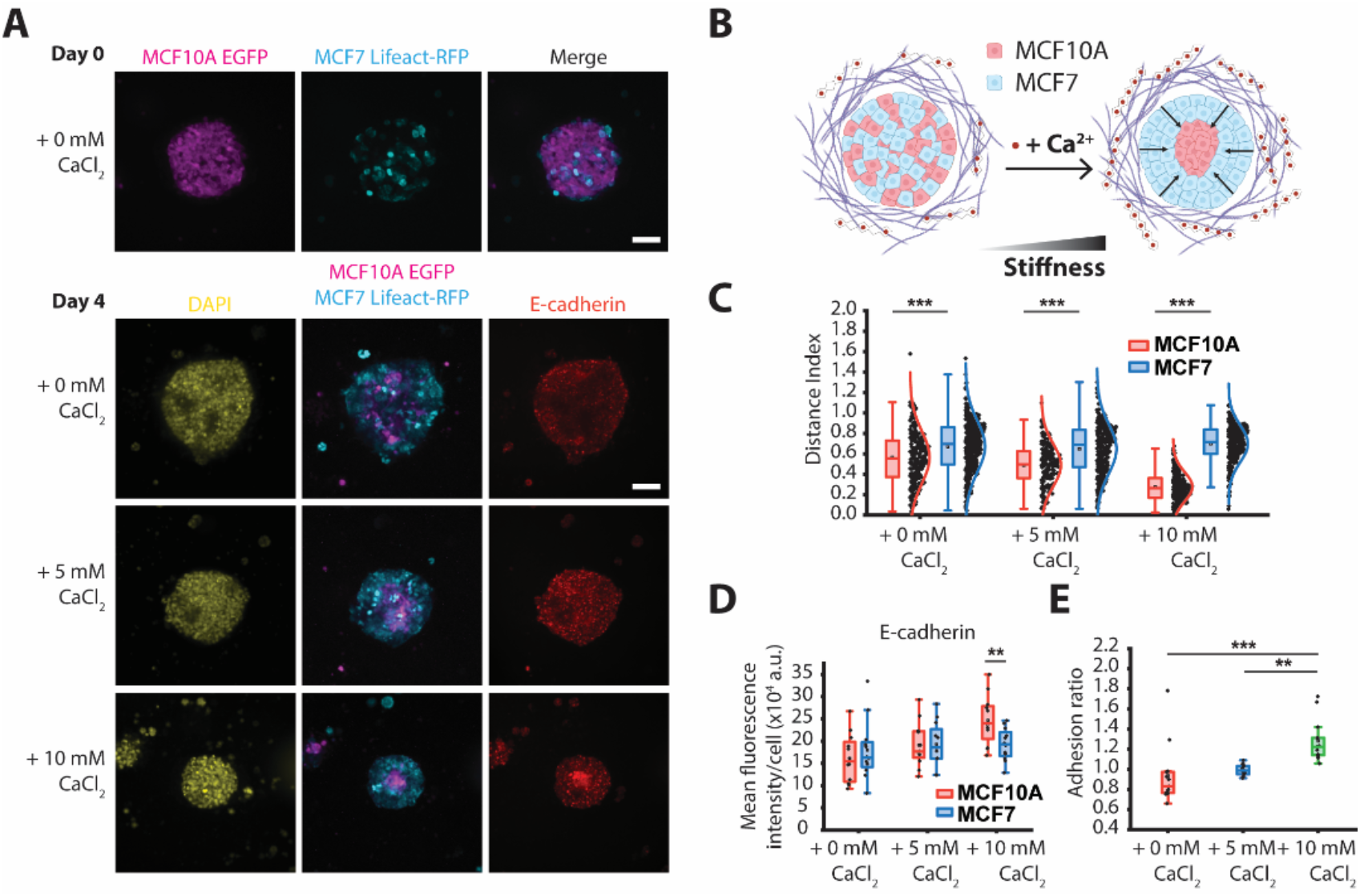
High confinement drives cell sorting in co-culture spheroids. **(A)** Fluorescence images of MCF10A and MCF7 co-culture spheroids encapsulated in collagen-alginate hydrogels and imaged on day 0 (day of encapsulation). 0, 5 or 10 mM CaCl_2_ was added on day 0, and the spheroids were imaged again on day 4. Spheroids were immunostained for E-cadherin and stained for DAPI. Fluorescence images show maximum projection of the z-slices. **(B)** Schematic depicting a MCF10A (red) and MCF7 (blue) co-culture spheroid embedded in a collagen-alginate hydrogel. Under high matrix confinement, MCF7 cells sort to the periphery and MCF10A cells form a cluster at the spheroid core. **(C)** Boxplot of distance index for MCF10A and MCF7 cells after 4 days of culture (*n =* 12 spheroids per condition). **(D)** Boxplot of E-cadherin fluorescence per cell for MCF10A and MCF7 co-culture spheroids on day 4 (*n =* 12 spheroids per condition). Only the median slice of the z-stack was utilized to calculate the distance index and E-cadherin fluorescence intensity. **(E)** Boxplot of the corresponding relative adhesion ratio of MCF10A and MCF7 co-culture spheroids. Scale bars are 90 μm. ** = *p* < 0.01, *** = *p* < 0.001.

Moreover, under high matrix confinement, mixed spheroids transitioned from a homogeneous adhesion state to a sorted state containing a defined core with high adhesion and a boundary compartment with low adhesion. These results suggest that physical confinement stress generated by the mechanical properties of the matrix was sensed by and transduced through cells, resulting in differential sorting. To confirm that the sorting behavior observed was not induced by the addition of calcium, we showed that the spheroids failed to sort when cultured in collagen gels with 10 mM CaCl_2_ (**Fig. S4**). In conditions of high matrix confinement, MCF7 cells sorted to the periphery, but lacked the ability to remodel the ECM [11]. Along with increasing matrix stiffness, we found that alginate crosslinking modified the ability of cells to remodel collagen fibers (**Fig. S5**), where there is a band of collagen fibers in the absence of additional calcium following 4 days of culture. In comparison, the accumulation of labeled collagen was less pronounced when alginate was crosslinked, indicating less remodeling. Together, these experiments indicate that, in our model system, cell sorting is driven by increased matrix confinement and is dependent on cell-cell adhesions.

### Tumor spheroid sorting depends on E-cadherin expression

When investigating the correlation between sorting and differential E-cadherin upregulation, we observed that spheroids consisting of MCF10A Lifeact-RFP and MCF10A EGFP failed to sort when subjected to high matrix confinement (**Fig. 3A**), consistent with our hypothesis that sorting is linked to differences in cell–cell junction strength and stability. In the absence of alginate, MCF10A spheroids collectively invaded into the surrounding matrix when cultured in collagen gels with 10 mM CaCl_2_ (**Fig. S6**). However, when embedded in collagen-alginate gels, MCF10A spheroids neither sorted nor invaded. Additionally, matrix confinement did not affect the size of the spheroids (**Figure S7**). We observed that spheroid size was not influenced by confinement conditions in unsorted 3D cultures, such as monoculture MCF10A spheroids. However, the size of MCF10A and MCF7 mixed spheroids significantly decreased as confinement increased and sorting occurred (**Fig. S3A**). In gels with high matrix stiffness, MCF10A E-cadherin levels were significantly lower, (**Fig. 3B**), and a similar result was found when MCF10A cells were co-cultured with invasive mesenchymal cells (**Fig. 1C**). Thus, the sorting behavior observed in mixed spheroids is likely attributed to the difference in E-cadherin expression of cancer cells and normal epithelial cells.

**Figure 3.**
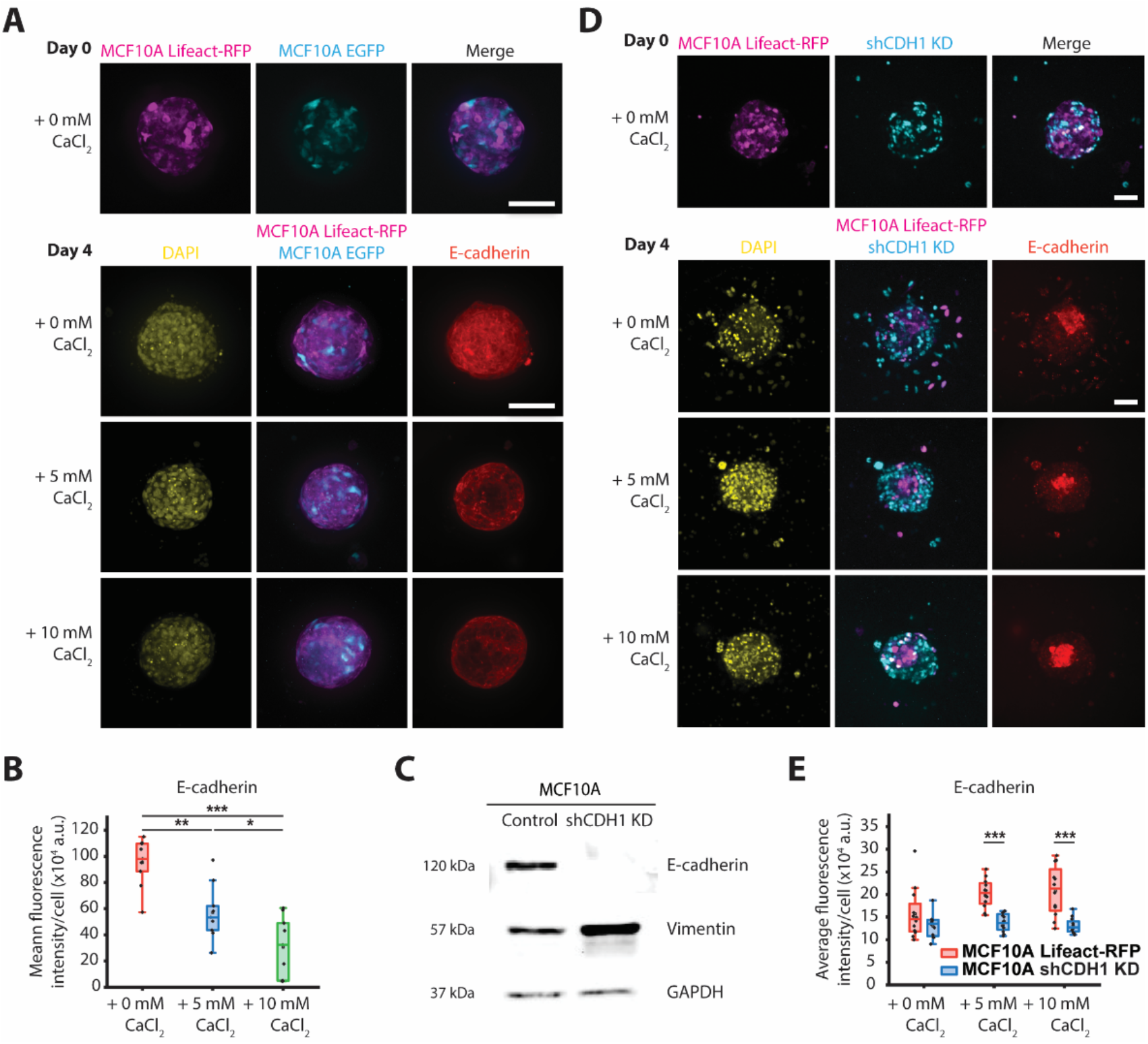
Tumor spheroid sorting depends on E-cadherin expression. **(A)** Representative fluorescence images of MCF10A Lifeact-RFP and EGFP co-culture spheroids encapsulated in collagen-alginate hydrogels and imaged on day 0 (day of encapsulation) and day 4 (day of fixation). Spheroids were immunostained for E-cadherin and stained for DAPI. **(B)** Boxplot of E-cadherin fluorescence per cell for MCF10A Lifeact-RFP and EGFP spheroids in hydrogels with 0, 5 or 10 mM CaCl_2_. **(C)** Western blot illustrating IPTG-induced shCDH1 KD in MCF10A cells and of vimentin expression in MCF10A scramble and shCDH1 KD cells. GAPDH was used as the loading control. **(D)** Representative fluorescence images of MCF10A Lifeact-RFP and shCDH1 KD mixed spheroids imaged on days 0 and 4. CaCl_2_ was added on day 0. Fluorescence images show maximum projection of the z-slices. **(E)** Boxplot of E-cadherin fluorescence per cell for MCF10A Lifeact-RFP and shCDH1 KD subpopulations in co-culture spheroids (*n =* 12 spheroids per condition). Only the median slice of the z-stack was utilized to calculate distance index and E-cadherin fluorescence intensity. Scale bars are 90 μm. * = *p* < 0.05, ** = *p* < 0.01, *** = *p* < 0.001.

Next, we asked if differences in E-cadherin expression regulated spheroid sorting. We knocked down E-cadherin (encoded by CDH1 gene) in MCF10A cells with inducible short hairpin RNA (shRNA) (**Fig. 3C**), as previously established [6,42], generating shCDH1 KD cells. Subsequently, MCF10A Lifeact-RFP and MCF10A shCDH1 KD co-culture spheroids were formed. The DAH proposes that strongly adhesive cells have a higher tissue surface tension and should preferentially adhere to each other and be enveloped by less adhesive cells. In agreement with the DAH, Lifeact-RFP cells clustered together at the core of the spheroid and were surrounded by layers of shCDH1 KD cells (**Fig. 3D**). In conditions of low confinement, individual cells detached from the spheroid and migrated into the surrounding matrix. When we calculated the distance index for the separate subpopulations within mixed spheroids, it revealed a distinct spatial separation between the two cell types under high confinement (**Fig. 3D**).

According to the DAH, tumors behave like fluids, and surface tension-like effects hold shCDH1 KD cells and non-invasive cancer cells within the spheroid boundary [17,59]. These cells are unable to break through compartment boundaries and invade into the surrounding matrix. shCDH1 KD cells in co-culture exhibit the same sorting behavior observed in MCF7 cells, showing that the sorting process is regulated by the difference in E-cadherin expression between the co-culture cell types (**Fig. 3E**).

### Sorted tumor spheroids unjam when matrix confinement is released

Having established that spheroid sorting is a result of increased matrix confinement, we then investigated whether reducing confinement will permit invasion. Addition of the calcium chelator EGTA relieves crosslinked alginate by binding to and removing calcium ions from the crosslinked hydrogel, lowering confinement. MCF10A Lifeact-RFP and MCF10A EGFP spheroids were used as a control and cultured with 10 mM CaCl_2_ (**Fig. 4A**). Then, after 4 days of culture, the spheroids were treated with 0 or 5 mM EGTA. Fluorescence imaging 2 days after treatment showed that EGTA-treated control spheroids did not migrate into the surrounding matrix (**Fig. 4A**), indicating that EGTA treatment and reduced confinement do not initiate invasion in unsorted spheroids.

**Figure 4.**
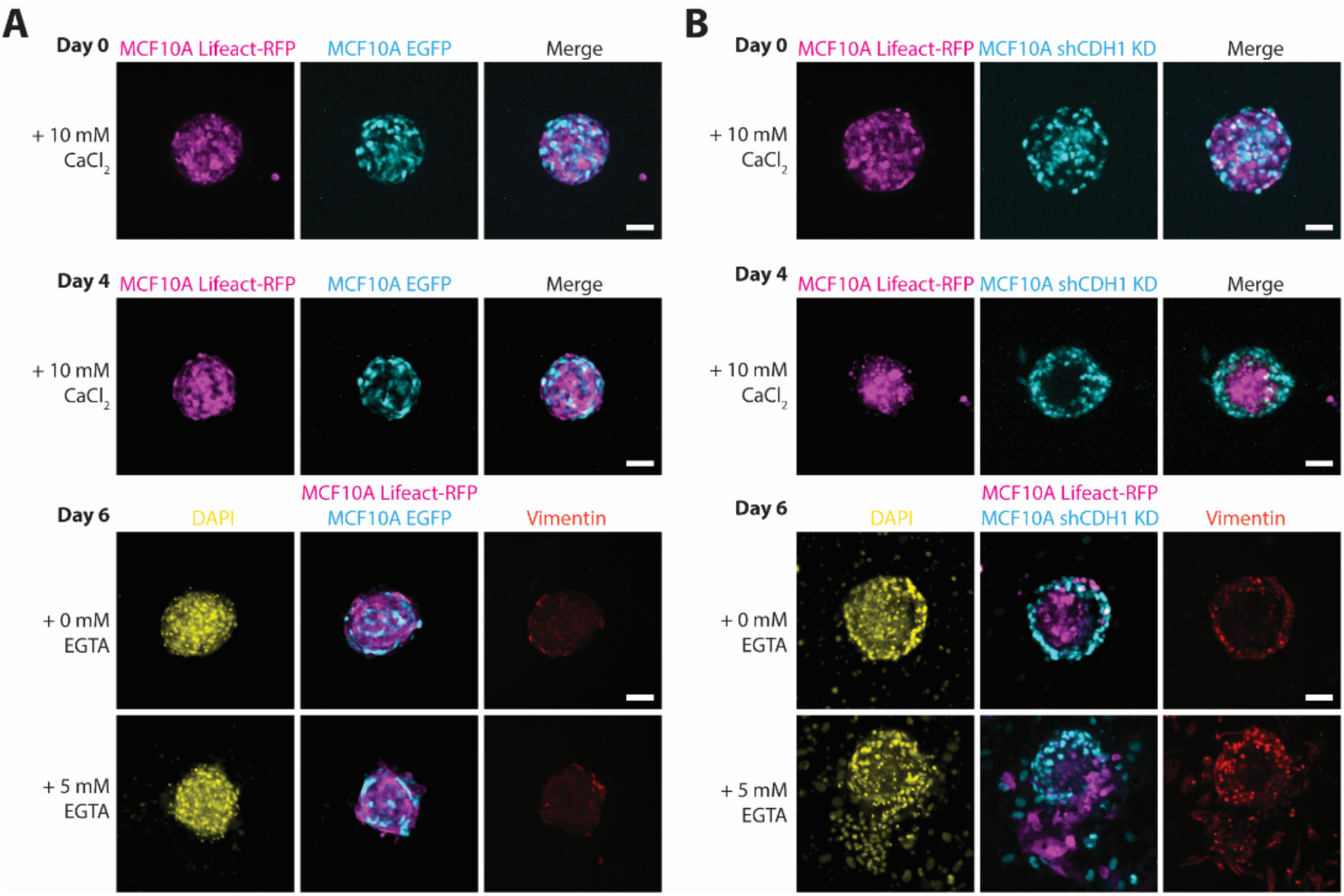
Sorted tumor spheroids unjam when matrix stiffness is lowered. **(A & B)** Representative fluorescence images of **(A)** MCF10A EGFP and **(B)** shCDH1 KD spheroids (co-cultured with MCF10A Lifeact-RFP cells) encapsulated in collagen-alginate hydrogels and imaged on day 0 (day of encapsulation), day 4, and day 6 (day of fixation). 0, 5 or 10 mM CaCl_2_ was added on day 0. On day 4, the spheroids were incubated with 0 or 5 mM EGTA for 1 h, after which the solution was replaced with fresh medium, and the spheroids were cultured until day 6. Spheroids were immunostained for vimentin and stained for DAPI. Fluorescence images show maximum projection of the z-slices. Scale bars are 90 μm.

Next, we explored whether lowering matrix confinement restores single-cell migration in MCF10A Lifeact-RFP and shCDH1 KD spheroids. We hypothesized that reversing the crosslinking reduces the confinement conditions responsible for spheroid sorting and for inhibiting cell migration into the matrix. Spheroids that were not treated with EGTA maintained a sorted state and were unable to invade. In EGTA-treated spheroids, MCF10A Lifeact-RFP cells emerged from the spheroid core by rupturing the surrounding shCDH1 KD cell layer (**Fig. 4B**). Downregulation of E-cadherin corresponded with upregulation of the mesenchymal marker vimentin (**Fig. 3C**), and vimentin expression in wildtype and shCDH1 KD cells was not influenced by confinement conditions (**Fig. S8**). As a benign epithelial cell line, MCF10A cells express low levels of vimentin (**Fig. 4A**), and as EMT is known to replace the keratin cytoskeleton with vimentin [17], the upregulation of vimentin observed in shCDH1 KD cells suggests a partial EMT. To rule out the effects of matrix viscoelasticity, we measured the loss tangent (the ratio of viscous to elastic effects) of our collagen-alginate composite gel using a rheometer (**Fig. S9**). We found that the loss tangent ranges from 0.05 to 0.12, but it does not exhibit a monotonic trend with respect to calcium concentration. In contrast, differential adhesion and the degree of cell sorting in co-cultured spheroids are correlated with calcium concentration. Thus, we believe the behaviors we observed are more closely related to the hydrogel stiffness than its viscoelasticity.

### Reducing matrix stiffness triggers burst-like migration in sorted spheroids

To examine the migration of sorted spheroids upon lowering matrix stiffness, we performed timelapse imaging for a 24-h period following EGTA treatment. Thus far, we have analyzed spheroid compartmentalization at different time scales, however time-lapse sequences are critical to capturing cell speed and trajectories. Since EGTA chelates calcium at a 1:1 ratio, we tested whether complete reversal of alginate crosslinking via the addition of 10 mM EGTA can further promote invasive behavior in co-culture spheroids. We generated two types of mixed spheroids: MCF10A Lifeact-RFP cells co-cultured with MCF10A EGFP or shCDH1 KD cells, and cultured the spheroids as previously described. On day 4, CaCl_2_ was removed from the medium, and the hydrogels were treated with 0 or 10 mM EGTA for 1 h prior to imaging. As expected, wildtype spheroids failed to sort or migrate regardless of EGTA treatment (**Fig. S10**), whereas untreated mixed shCDH1 KD spheroids remained in a sorted state over the 24-h period (**Movie S2 and Fig. 5A**, top). At the spheroid core, where cell density is high, MCF10A Lifeact-RFP cells were jammed, whereas the surrounding shCDH1 KD cells were in a more fluid-like state, confined by the barrier of high matrix confinement (**Fig. S11 and Fig. 5B**, top). In sorted spheroids, EGTA treatment reduced matrix confinement, resulting in the dissemination of both cell types (**Fig. 5A and Fig. 5B**, bottom). Cells that were sorted to the spheroid core rapidly migrated outwards into the matrix with high motility (**Movies S2 and S3**).

**Figure 5.**
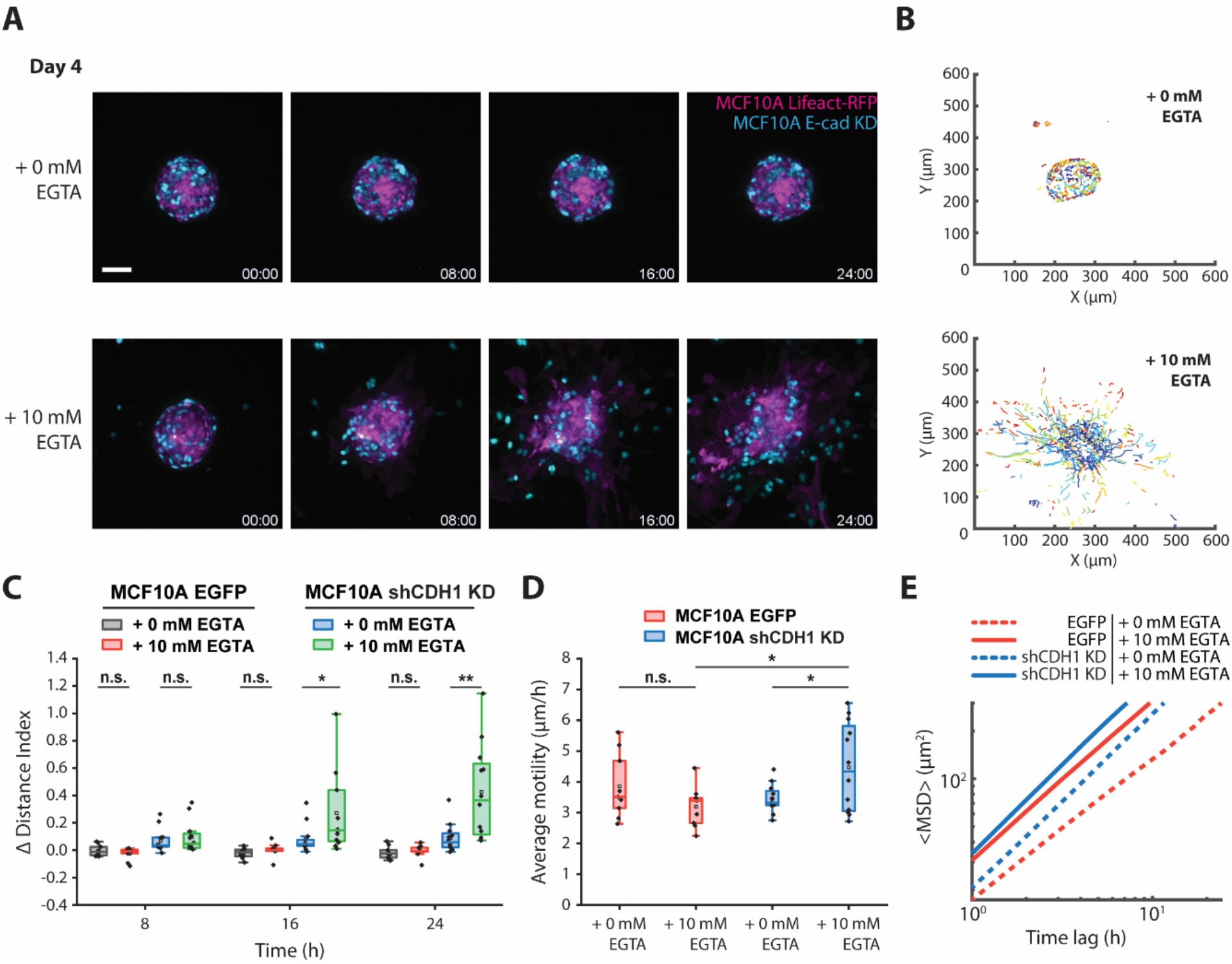
The reversal of alginate crosslinking stimulates burst-like migration in sorted spheroids. **(A)** Timelapse of MCF10A Lifeact-RFP and shCDH1 co-culture spheroids over a 24-h period, starting on day 4 after incubation with 0 or 10 mM EGTA for 1 h, after which the solution was replaced with fresh medium prior to time-lapse imaging. Fluorescence images show maximum projection of the z-slices. **(B)** Corresponding tracked trajectories of MCF10A shCDH1 KD cells in spheroids treated with 0 mM EGTA (top) and 10 mM EGTA (bottom). **(C)** Boxplot of distance index for MCF10A EGFP or shCDH1 KD cells in co-culture spheroids (with MCF10A Lifeact-RFP cells) analyzed at 8-hour intervals over 24 h. A positive change in distance index indicates cell motion away from the spheroid core. Only the median slice of the z-stack was utilized to calculate distance index. **(D)** Boxplot of average cell motility for MCF10A EGFP or shCDH1 KD cells (*n =* 9–12 spheroids per condition). **(E)** MSDs for MCF10A EGFP or shCDH1 KD cells plotted over a 24-h period. Each line represents the mean MSD for *n =* 12 spheroids. The plots are displayed in log-log scale. Z-stacks were used to calculate cell motility and MSD. Scale bars are 90 μm.

The results presented thus far led us to hypothesize that the jamming—unjamming transition in 3D can be manipulated by modulating matrix confinement. To statistically quantify the spatial organization of spheroids over time, we tracked individual MCF10A EGFP or MCF10A shCDH1 KD cells within the two types of mixed spheroids and calculated the change in distance index compared to the initial time point. The average distance travelled by shCDH1 KD cells away from the spheroid core in treated spheroids was significantly higher than in non-treated spheroids (**Fig. 5C**). Corresponding to the change in distance index (quantified as the difference between the mean distance index of cells for a given spheroid at a specified time point and the cells’ initial mean distance index), EGTA treatment triggered and accelerated the migration of shCDH1 KD cells, whereas wildtype MCF10A cell motility was not impacted (**Fig. 5D**). Computation of the MSD revealed that EGTA treatment increased diffusivity for both cell types, however shCDH1 KD cells were more diffusive than their wildtype counterparts under the same conditions (**Fig. 5E**).

### In theory and experiment, matrix confinement governs spheroid sorting and invasion

In order to understand the impact of matrix confinement on cellular invasion, we utilized a SPV model [45,46], which mimics the dynamics of a heterogeneous tissue comprised of two different types of cells. Our simulations produced a well-sorted tissue, a result of setting differential adhesions parameters [59,60] at cell-cell junctions (**Fig. S12**). Using the sorted tissue as a starting point, we manipulated the cell-medium contact tensions (*τ)* to study changes in cell sorting and invasion behaviors.

**Fig. 6A** presents representative snapshots illustrating cellular dispersion under conditions of high (left), medium (middle) and low (right) confinement stress (*σ*_*n*_). These computational results suggest that cells in a well-sorted state manage to breach their boundaries and invade the ECM when confinement is diminished. In order to quantify this cellular dispersion of cells, we calculated a sorting index, defined as *I* = 1 − *N*_*cluster*_/*N*, where *N*_*cluster*_ is the count of isolated cell clusters and *N* is the total cell count for each cell type. In instances where the cells are fully sorted, each cell type would form a single cluster, resulting in a sorting index of *I* ≈ 1. If the cells are dispersed, they form a mixed state, and each cell forms an isolated cluster unconnected from others, making the sorting index 0. The sorting index *I*, as a function of *σ*_*n*_, is shown in **Fig. 6B**. We can classify the behavior of the cell dispersion into three distinct phases. On the left of the white dashed line, the tissue experiences high confinement. Both A and B cells remain sorted with sorting index *I* ≈ 1, corresponding to the left snapshot of **Fig. 6A** (marked by a star). The middle region, located between the white and black dashed lines, shows that the outer-layered red B cells have dispersed while the A cells stayed sorted, corresponding to the middle snapshot in **Fig. 6A** (marked by a circle). On the right of the black dashed line, the confinement has essentially vanished. The B cells have completely dispersed, and A cells begin to invade into the ECM, corresponding to the right snapshot in **Fig. 6A** (marked by a square).

**Figure 6.**
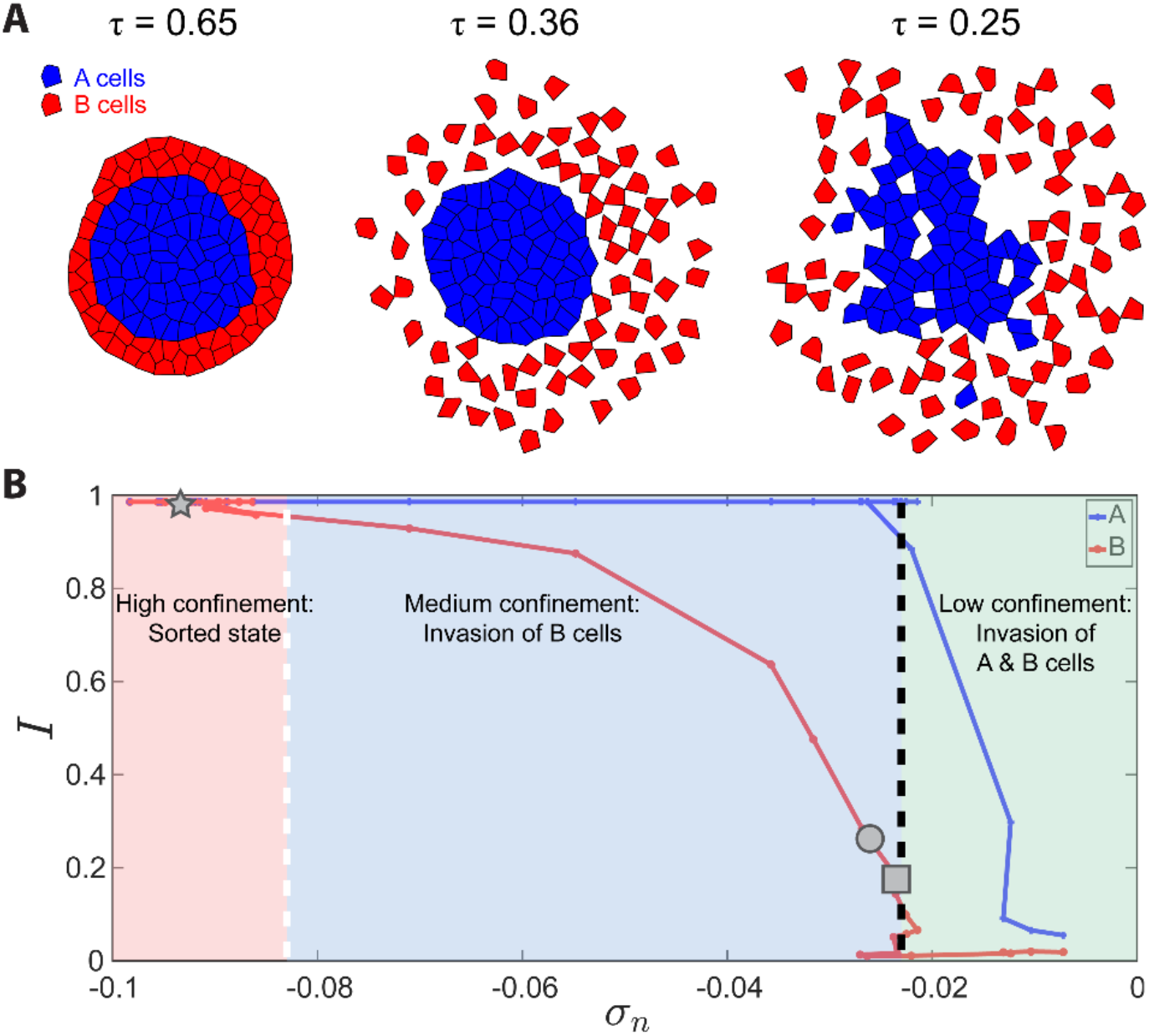
SPV simulations portraying a heterogeneous tissue with two cell types, blue A cells and red B cells. **(A)** Representative snapshots of cell invasion into the ECM under varying confinement levels. Left: high confinement prevents dispersion. Middle: dispersion of the outer-layered B cells. Right: both A and B cells invade under low confinement. The three snapshots are marked as star, circle, and square in panel **(B). (B)** Cell sorting index *I* is plotted as a function of compressive stress *σ*_*n*_. By downregulating *σ*_*n*_, the cells could disperse into the ECM and the sorting index decreases. The dashed white and black lines denote the commencement of dispersion for B and A cells respectively. The invasion behavior bifurcates into three distinct regimes: (1) Red region to the left of the white dashed line indicates high confinement and a sorted tissue. (2) Blue region between white and black dashed lines signifies dispersed B cells and sorted A cells. (3) Green region to the right of the black dashed line represents vanishing compressive stress and unsorted A and B cells.

Taken together, our results corroborate a model in which matrix confinement promotes spheroid sorting in an adhesion-dependent manner and subsequently affects cellular unjamming processes (**Fig. 7**). Starting from a mixture of wildtype and shCDH1 KD spheroids encapsulated in collagen-alginate gels, the addition of 10 mM CaCl_2_ crosslinked the hydrogel, increasing matrix confinement, which caused the spheroids to sort. The ability of cells to invade the surrounding matrix hinges on the competition between the forces generated by the cells and the resistance exerted by the matrix. Crosslinking the hydrogel amplifies its yield stress which hinders spheroid volume expansion [69]. At the spheroid core where cell density is high, the adhesive wildtype cells cannot overcome the solid stress to intercalate positions with neighboring cells, leading to a jammed state [70]. Conversely, shCDH1 KD cells at the spheroid periphery are not jammed, however the cells lack sufficient energy to escape their compartment boundary. By reversing the crosslinking, the resistance to cell motion is removed, and the cells flow in an unjamming transition, reminiscent of the release of solid stress and elastic energy when the mechanical confining structure of an excised tumor is disrupted [71]. 5 mM EGTA treatment partially reversed the crosslinking and correspondingly reduced ECM confinement. As a result, the compacted wildtype cells escaped from the spheroid core as a strand and broke through the surrounding layers of shCDH1 KD cells. When the crosslinking was completely reversed by the addition of 10 mM EGTA, the cells displayed collective and super-diffusive motion as they were propelled into the matrix with high velocity (**Movie S1**).

**Figure 7.**
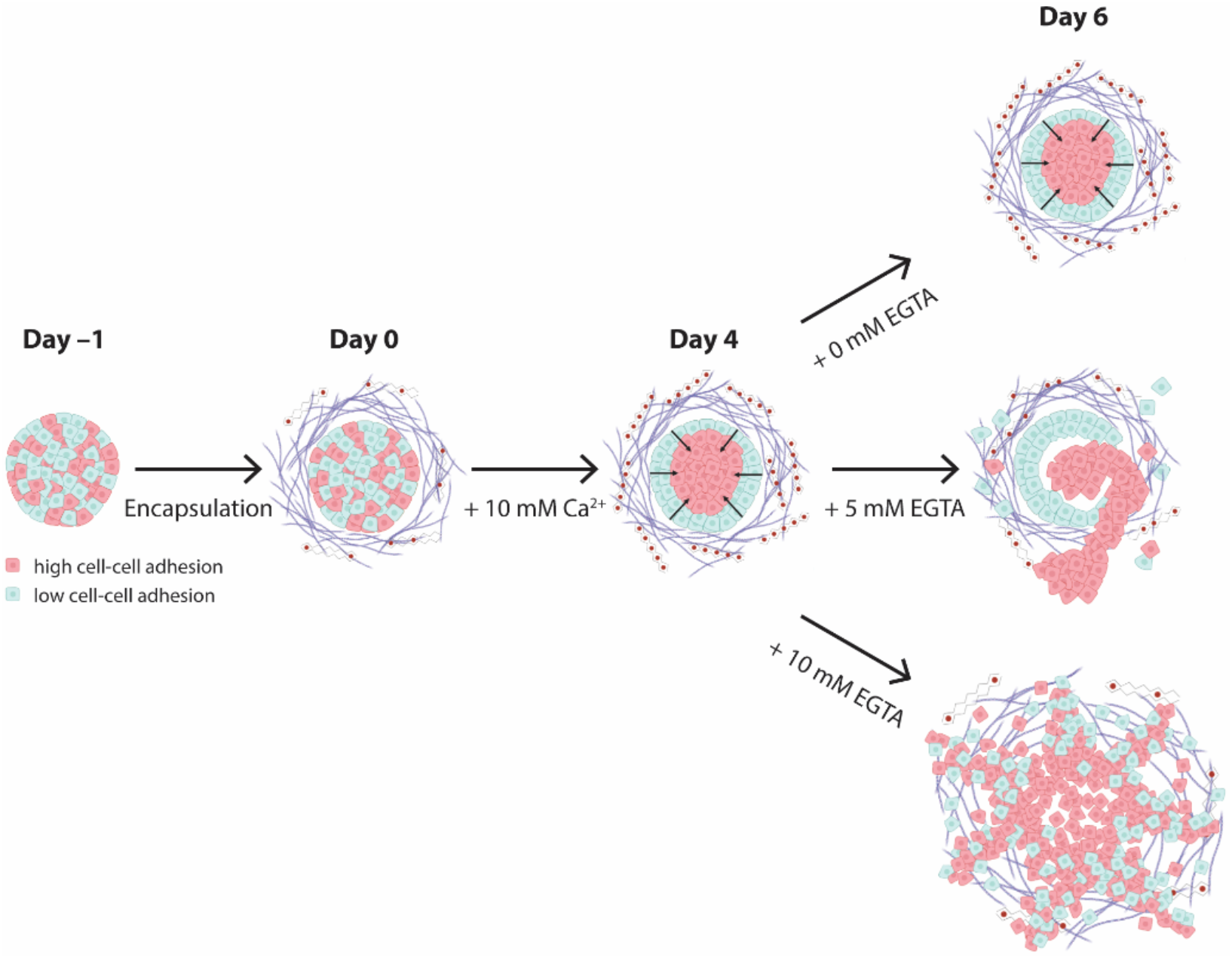
Tumor spheroid sorting and burst-like migration in a composite hydrogel with tunable stiffness. Summary depicting a spheroid generated from a mixture of cells with high cell-cell adhesion (red) and cells with low cell-cell adhesion (blue) on day –1, encapsulation of the spheroid in a collagen-alginate hydrogel on day 0 and sorting under high-confinement ECM on day 4. The rightmost panel depicts, on day 6, the spheroid sustaining the sorted state after 0 mM EGTA treatment (top), a strand-like cluster emerging from the spheroid core after 5 mM EGTA treatment (middle) and burst-like unjamming after 10 mM EGTA treatment (bottom).

## Discussion

In the present study, we show that tuning matrix confinement regulates 3D spheroid sorting and collective cell migration into the ECM. ECM stiffness is known to play a significant role in cancer spread and metastasis in the body [72–74], and we first establish that single-cell cancer migration is restricted by matrix stiffness (**Fig. 1**). We then demonstrate the sorting of non-invasive cancer cells and normal breast epithelial cells in co-culture spheroids (**Fig. 2**). The DAH assumes that sorting is a result of cells rearranging to minimize interfacial tension, which is directly proportional to differences in cell-cell adhesion [17,23,59], and as a result, spheroid sorting achieves an equilibrium thermodynamic state [59]. In hydrogels with minimal crosslinking, spheroids grow over four days of culture without exhibiting sorting behavior. However, spheroids were observed to sort in conditions of high matrix confinement (**Figure 2A**). In monoculture spheroids of benign breast epithelial cells, E-cadherin expression is reduced in response to high matrix confinement (**Fig. 3A**). One possibility is that in the absence of mesenchymal cells to induce sorting and secrete enzymes to remodel the matrix, normal cells undergo a partial EMT to promote invasion [75]. Previous studies have established that the unjamming transition is distinct from EMT in 2D cultures [76,77]. Benign cells may adapt to confined conditions by adjusting the properties of the cellular collective, however a stiff matrix environment presents a barrier that the cells are unable to overcome.

When hydrogels are crosslinked, they create a barrier that hinders cell movement into the matrix, leading to cell jamming. Conditions of high confinement enhance cell sorting within a co-culture model through the regulation of cell adhesion properties, which amplifies the differential adhesiveness between distinct cell populations. This results in a highly pronounced sorting of cells based on their adhesive properties within the co-culture model. Conversely, it is possible that cells exhibit a reduced tendency to sort in low confinement conditions because their adhesive interactions with neighboring cells are weak, which is a potential barrier to cell sorting. Stably downregulating E-cadherin identifies differences in intercellular adhesion as the primary driver of cell sorting (**Fig. 3B**). A possible explanation is that spheroid sorting creates a high local cell density in the core, where cells enhance their adhesions, reduce their volume, and display jammed behavior. In this state, the cells are restricted by adhesion, leading to increased tension along cell-cell interfaces [23,78,79].

One open question is how stress fluctuations driven by nearby cell division impact cell motility and structural rearrangements in the dense spheroid core [23]. Since less adhesive mesenchymal cells are relegated to the outer edges of the spheroid, core cells require energy to escape the boundary constraints imposed by both their neighbors and the surrounding matrix. Cell sorting may increase internal spheroid pressure, and thus, reducing matrix stiffness would lead to a pressure differential between the spheroid and the surrounding matrix. Raghuraman *et al*. recorded burst-like cellular motion in soft collagen matrices (0.5 mg/ml) and attributed it to the pressure increase within the cancer aggregate due to cellular swelling [80]. In this study, we provide evidence that manipulating matrix confinement directly impacts 3D spheroid sorting, as well as individual and collective cell invasion. In comparison to single-cell cancer migration, burst-like migration provides a potential outlet for cells confined by the spheroid boundary and for tumor cells to rapidly escape into the surrounding matrix. Further studies are required to test this model and explore the impact of stress fluctuations from nearby cell division on cell motility and structural changes in different regions of 3D spheroids.

Our hydrogel system closely replicates the tumor microenvironment and facilitates the study of cell-cell and cell-matrix interactions within a 3D matrix, which is more physiologically relevant compared to 2D culture systems. The ability to adjust and reverse the crosslinking, and thus the stiffness of the hydrogel matrix, by tuning the calcium concentration and adding a calcium chelator, enables dynamic studies of cell behavior in response to changes in matrix confinement over time. Our results show how matrix confinement leads to changes in surface tension at the cell-ECM interface, influencing cellular sorting and spheroid dispersion. However, an unresolved question is whether the sorting process is kinetically hindered under conditions of low confinement. This hindrance could arise from several factors: the physical constraints of expanding spheroids, which result in lower cell density; the early departure of less adhesive cells; or a diminished inclination for cells to sort due to weaker adhesive interactions with neighboring cells when matrix confinement is low. These points highlight the need for further research into the kinetic barriers to cell sorting and the impact of confinement on cellular organization and migration.

To better understand how cells in spheroids respond to mechanical stimuli, analyzing expression levels of mechanotransduction markers is crucial in understanding how cells perceive and respond to mechanical signals [81]. Exploring nuclear mechanosensing in this context would be particularly interesting, as it could provide deeper insights into how cells respond to mechanical stimuli at the nuclear level [82]. This can shed light on how mechanical forces affect crucial processes such as gene expression and DNA repair.

In tumor development, cancer cells are confined by compartment boundaries until a late stage [17,83]. Primary tumors are encapsulated and confined by a basement membrane, and as the tumor cells proliferate, the tumor experiences high confinement stress and may be driven to sort according to the DAH. In the case of tumor cell sorting, mesenchymal cells will preferentially sort to the spheroid boundary, and a reduction in ECM confinement can instigate an unjamming transition, leading to a rapid spread of tumor cells. Our work presents new insights into how matrix mechanical properties impact the mechanisms of collective cell motion within primary tumors and cancer migration to distant metastatic sites, both as individual cells and as cellular aggregates.

## Acknowledgements

The authors thank the Liu lab, particularly Sung-Won Hwang and Hossein Moghimianavval, and Ovijit Chaudhuri (Stanford), for helpful discussions.

## Author contributions

Conceptualization: G.C. and A.P.L. Methodology: G.C., X.L., S.J.C., N.C.R., and K.K. Investigation: G.C., S.-S.L., S.J.C., N.C.R., and K.K. Visualization: G.C., X.L., S.J.C., and N.C.R. Computational modeling: X.L. and D.B. Supervision: D.B. and A.P.L. Writing—original draft: G.C. and X.L. Writing—review & editing: G.C., S.-S.L., D.B., S.J.C., N.C.R., and A.P.L.

## Data availability

The data required to reproduce these findings are available upon request.

