## Supplementary Materials for "Matrix confinement modulates 3D spheroid sorting and burst-like collective migration"

**This PDF file includes:**

Figs. S1 to S12

Legends for movies S1-S3

**Other Supplementary Material for this manuscript includes the following:**

Movies S1-S3

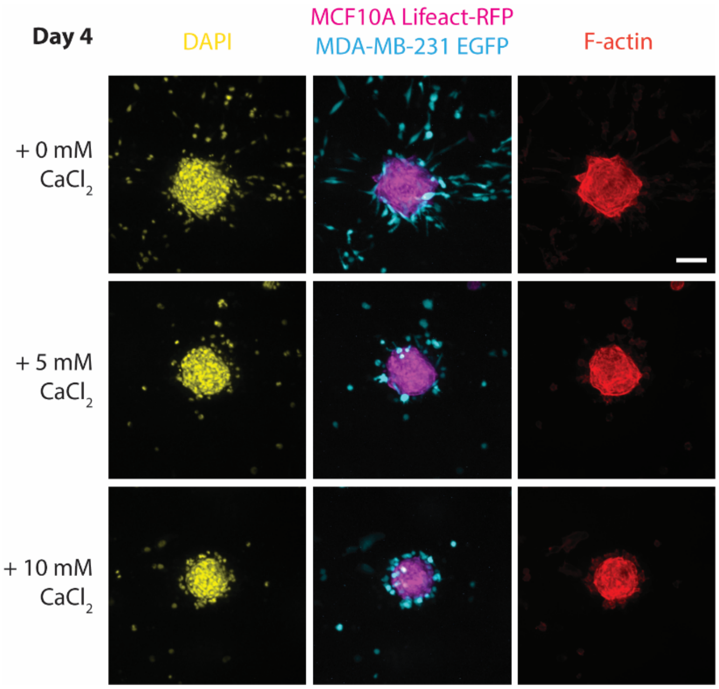


**Figure S1. High confinement inhibits cancer cell migration in collagen-alginate hydrogels.** Representative fluorescence images of MCF10A Lifeact-RFP and MDA-MB-231 EGFP co-culture spheroids embedded in collagen-alginate hydrogels with different concentrations of $CaCl_2$ and stained for DAPI and F-actin after 4 days of culture. Scale bar: 90 µm.

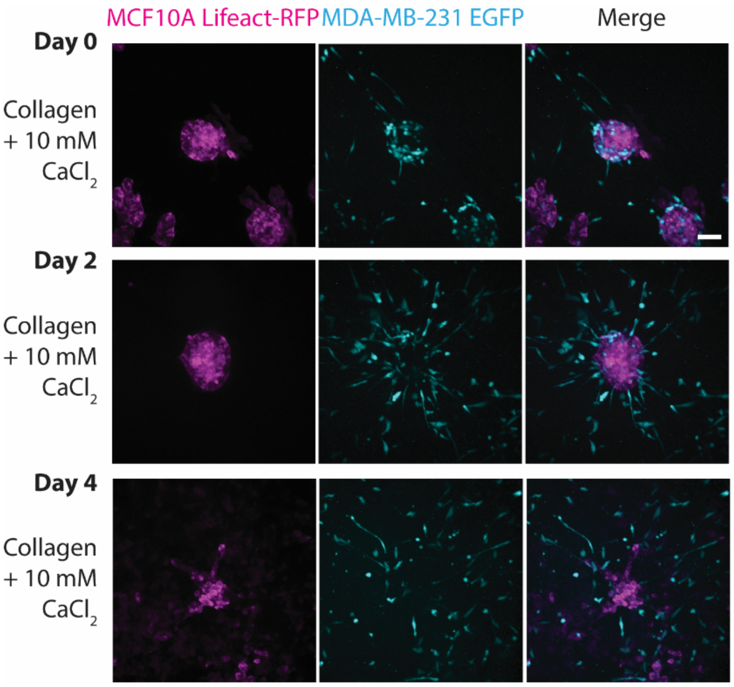


**Figure S2. MCF10A and MDA-MB-231 mixed spheroids cultured in collagen hydrogels invade into the surrounding matrix.** Representative fluorescence images of MCF10A Lifeact-RFP and MDA-MB-231 EGFP co-culture spheroids embedded in collagen hydrogels with different concentrations of $CaCl_2$ and imaged at days 0, 2 and 4. On day 0, the spheroids were encapsulated and 10 mM $CaCl_2$ was added. Scale bar: 90 µm.

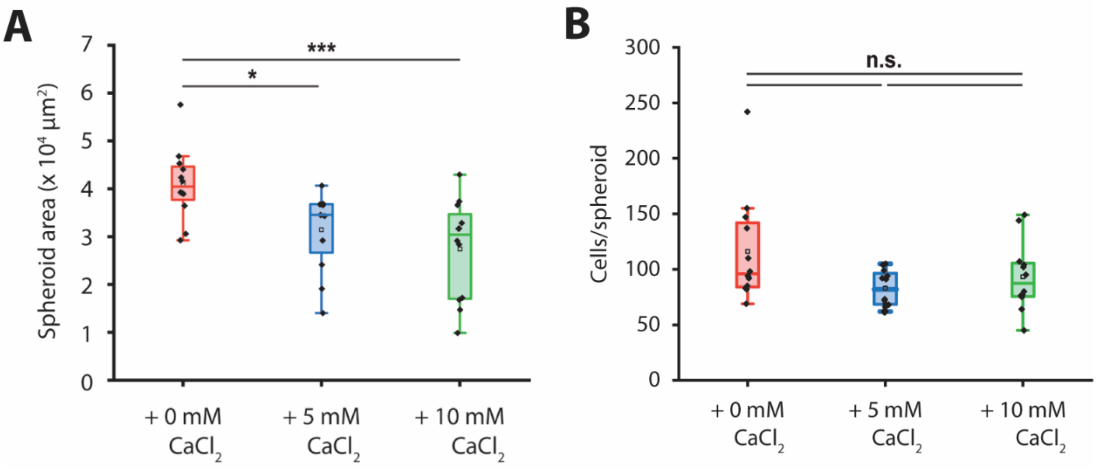


**Figure S3. The size of MCF10A and MCF7 mixed spheroids decreases with increasing matrix stiffness. (A)** Boxplot of spheroid area for MCF10A and MCF7 co-culture spheroids in collagen-alginate hydrogels after 4 days of culture with 0, 5 or 10 mM $CaCl_2$. **(B)** Boxplot showing the number of cells per spheroid. The number of DAPI-stained nuclei was automatically counted in Fiji. * = $p < 0.05$, *** = $p < 0.001$.

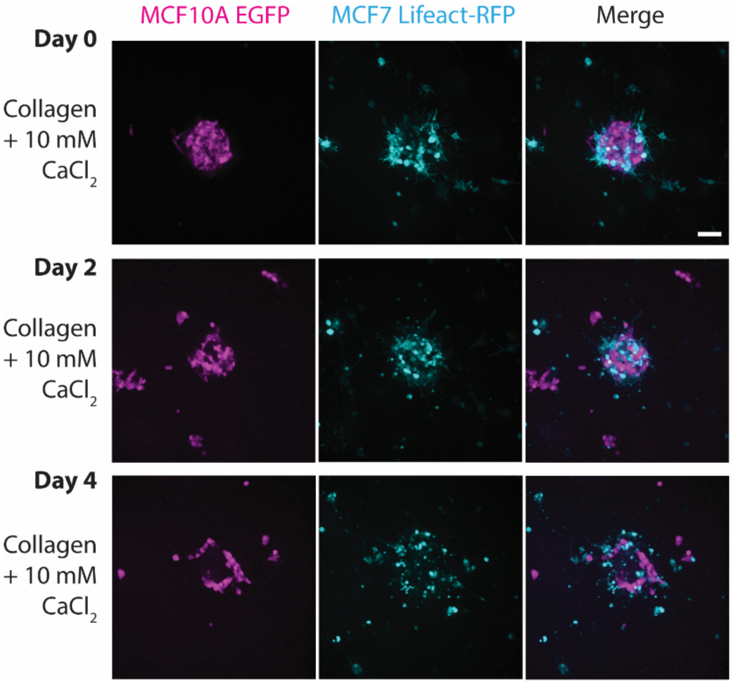


**Figure S4. MCF10A and MCF7 co-culture spheroids fail to sort in collagen hydrogels.** Representative fluorescence images of MCF10A EGFP and MCF7 Lifeact-RFP co-culture spheroids embedded in collagen hydrogels with different concentrations of $CaCl_2$ and imaged at days 0, 2 and 4. On day 0, the spheroids were encapsulated and 10 mM $CaCl_2$ was added. Scale bar: 90 µm.

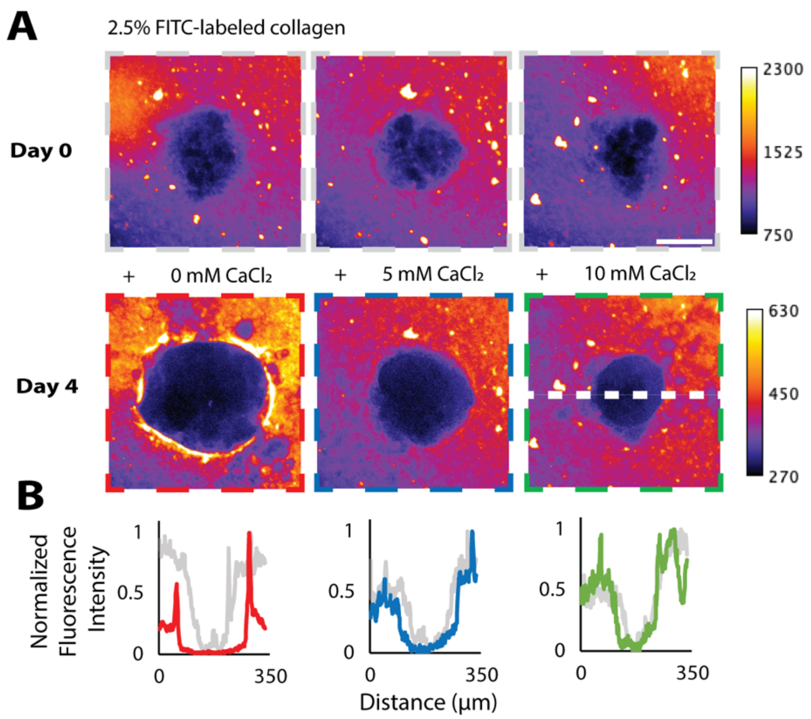


**Figure S5. Local ECM deformation minimized in calcium-crosslinked collagen-alginate hydrogels. (A)** Representative maximum projection images ofMCF10A and MCF7 co-culture spheroids suspended in 2.5% FITC-labeled collagen and 0.25% alginate hydrogel. Local collagen network was visualized before (top) and 4 days after (bottom) addition of $CaCl_2$ to crosslink alginate and increase hydrogel stiffness. White dashed line represents line scans used to quantify fluorescence intensity. Scale bar: 100 µm. **(B)** Fluorescence intensity line scans of FITC-labeled collagen. Lines were drawn across the spheroid, as shown in panel A. Line colors correspond with the color of the dashed boxes in panel A.

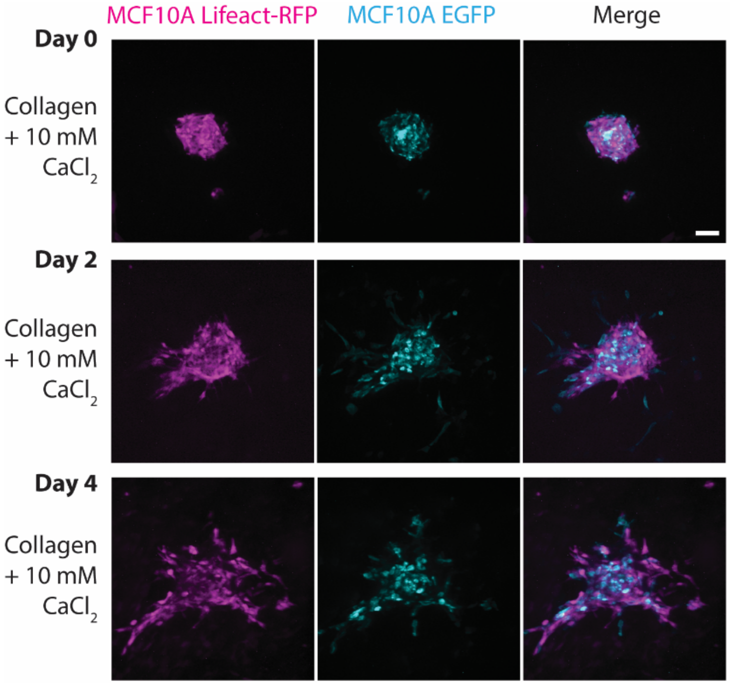


**Figure S6. MCF10A spheroids cultured in collagen hydrogels invade into the surrounding matrix.** Representative fluorescence images of MCF10A Lifeact-RFP and EGFP spheroids embedded in collagen hydrogels and imaged at days 0, 2 and 4. On day 0, the spheroids were encapsulated and 10 mM $CaCl_2$ was added. Scale bar: 90 µm.

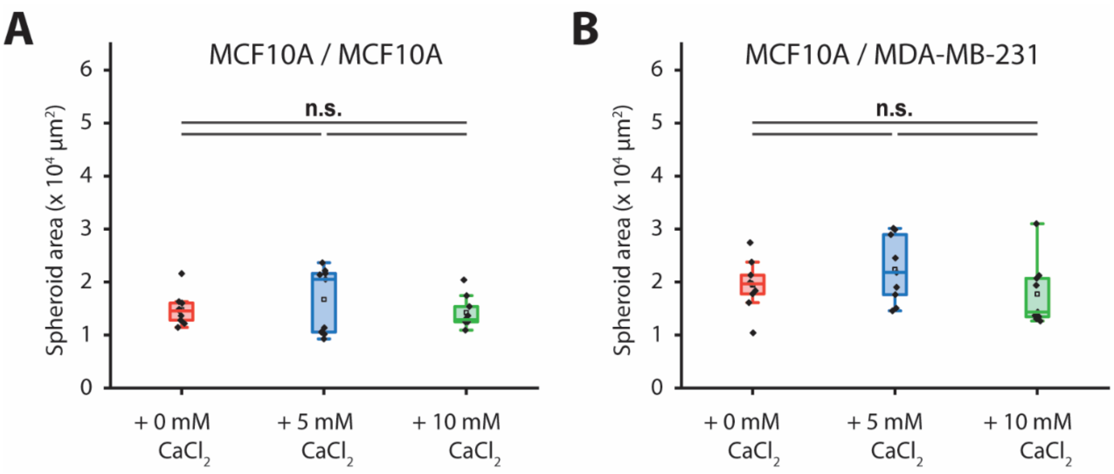


**Figure S7. The level of confinement does not influence the size of MCF10A monoculture spheroids. (A)** Boxplot of spheroid area for MCF10A monoculture spheroids in collagen-alginate hydrogels after 4 days of culture with 0, 5 or 10 mM $CaCl_2$. **(B)** Boxplot of spheroid area for MCF10A and MDA-MB-231 co-culture spheroids in collagen-alginate hydrogels after 4 days of culture with 0, 5 or 10 mM $CaCl_2$.

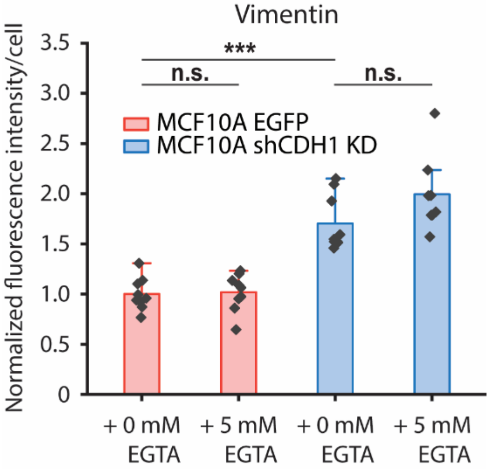


**Figure S8. Vimentin expression was not influenced by changes in matrix stiffness.** Vimentin fluorescence per cell for MCF10A EGFP or shCDH1 KD cells in co-culture spheroids (with MCF10A Lifeact-RFP cells). On day 0, the spheroids were encapsulated in collagen-alginate hydrogels and 10 mM $CaCl_2$ was added. The samples were subsequently treated with 0 or 5 mM EGTA on day 4 and stained for DAPI and vimentin on day 6. *** = $p < 0.001$.

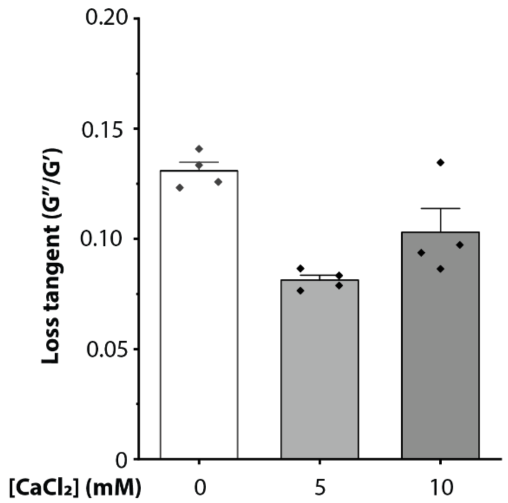


**Figure S9. Loss tangent of collagen-alginate hydrogel with different added calcium concentrations.** Viscoelastic measurements were conducted at 1 rad/s and 1% strain for 1,000 seconds, and measurements were taken every 10 seconds. Four independent samples were characterized. Error bars denote S.E.

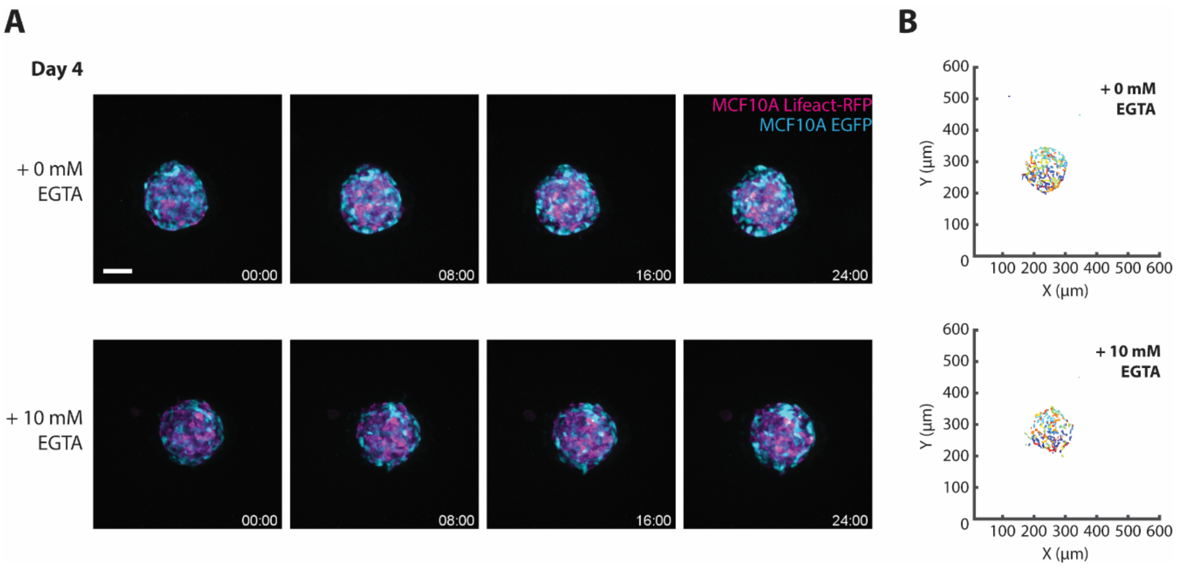


**Figure S10. Wildtype spheroids do not sort or migrate regardless of EGTA treatment. (A)** Timelapse of MCF10A Lifeact-RFP and EGFP co-culture spheroids over a 24 h period, starting on day 4 after incubation with 0 or 10 mM EGTA for 1 h, after which the solution was replaced with fresh medium prior to time-lapse imaging. **(B)** Corresponding tracked trajectories of MCF10A EGFP cells in spheroids treated with 0 mM EGTA (top) and 10 mM EGTA (bottom). Scale bar: 90 μm.

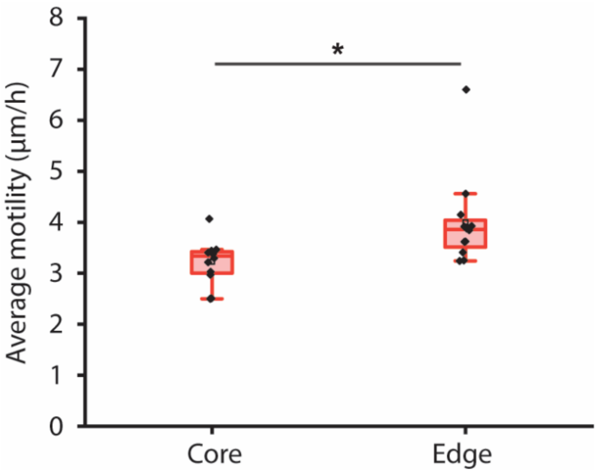


**Figure S11. Sorted MCF10A Lifeact-RFP and shCDH1 KD spheroids exhibit higher cell motility at the spheroid edge.** Boxplot of average cell motility in the core vs. the periphery for MCF10A Lifeact-RFP and shCDH1 mixed spheroids in collagen-alginate hydrogels after 4 days of culture with 10 mM $CaCl_2$. * = $p < 0.05$.

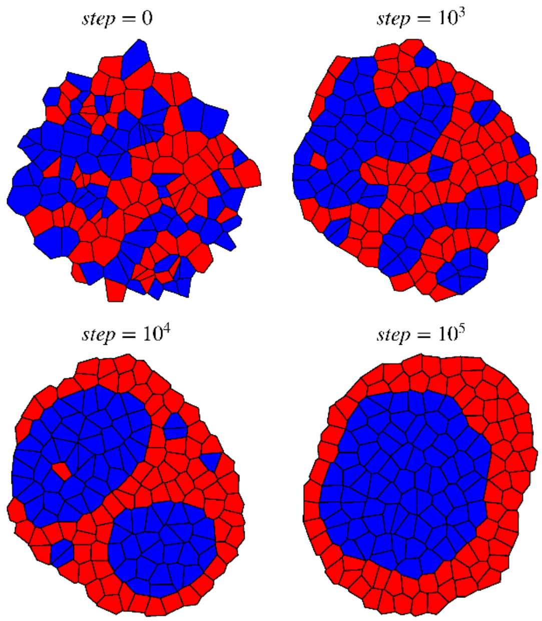


**Figure S12. Segregation of a heterogeneous tissue with two cell types, blue A cells and red B cells under differential adhesions.** Differential adhesion values are set to be $\tau_{AB} > \frac{\tau_{AA}+\tau_{BB}}{2}$ to achieve a sorted state.

**Supplementary movie legends**

**Movie S1.** Representative z-stack of a MCF10A EGFP and MCF7 Lifeact-RFP mixed spheroid cultured in collagen-alginate hydrogel with 10 mM $CaCl_2$ and imaged after 4 days of culture. Scale bar is 90 μm.

**Movie S2.** Representative movie of a MCF10A Lifeact-RFP and shCDH1 KD mixed spheroid cultured in collagen-alginate hydrogel with 10 mM $CaCl_2$ and imaged over a 24-h period after 0 mM (left) and 10 mM (right) EGTA treatment. Scale bar is 90 μm.

**Movie S3.** Representative z-stack of a MCF10A Lifeact-RFP and shCDH1 KD mixed spheroid cultured in collagen-alginate hydrogel with 10 mM $CaCl_2$ and imaged 14 h after 10 mM EGTA treatment. Scale bar is 90 μm.
